# Stochastic dynamics of an epidemics with recurrent spillovers from an endemic reservoir

**DOI:** 10.1101/213579

**Authors:** Marina Voinson, Alexandra Alvergne, Sylvain Billiard, Charline Smadi

## Abstract

Most emerging human infectious diseases have an animal origin. Yet, while zoonotic diseases originate from a primary reservoir, most theoretical studies have principally focused on single-host processes, either exclusively humans or exclusively animals, without considering the importance of animal to human transmission for understanding the dynamics of emerging infectious diseases. Here we aim to investigate the importance of spillover transmission for explaining the number and the size of outbreaks. We propose a simple stochastic Susceptible-Infected-Recovered model with a recurrent infection of an incidental host from a reservoir (e.g. humans by a zoonotic species), considering two modes of transmission, (1) animal-to-human and (2) human-to-human. The model assumes that (i) epidemiological processes are faster than other processes such as demographics or pathogen evolution and (ii) that an epidemic occurs until there are no susceptible individuals left. The results show that during an epidemic, even when the pathogens are barely contagious, multiple outbreaks are observed due to spillover transmission. Overall, the findings demonstrate that the only consideration of direct transmission between individuals is not sufficient to explain the dynamics of zoonotic pathogens in an incidental host.

## 1. Introduction

Recent decades have seen a surge of emerging infectious diseases (EIDs), with up to forty new diseases recorded since the 1970s [1]. Sixty percent of emerging human infectious diseases have an animal origin [1, 2]. The World Health Organization defines zoonotic pathogens as “pathogens that are naturally transmitted to humans via vertebrate animals”. The epidemics caused by EIDs impact the societal and economical equilibria of countries by increasing unexpected deaths, the need for health care infrastructures and by interfering with travel [3]. Moreover, the risk of EIDs being transmitted to humans from wildlife is increasing because of the recent growth and geographic expansion of human populations, climate change and deforestation, which all increase the number of contacts between humans and potential new pathogens [1, 4, 5]. Given this, it is crucially important to understand how infections spread from reservoir, i.e. by spillover transmission.

There is numerous empirical evidence that the epidemiological dynamics of infectious diseases is highly dependent on the spillover transmission from the reservoir (for the reservoir definition see Table 1). The start of an outbreak is promoted by a primary contact between the reservoir and the incidental host (i.e. host that becomes infected but is not part of the reservoir) leading to the potential transmission of the infection in the host population. Moreover, multiple outbreaks are commonly observed during an epidemic of zoonotic pathogens in human population, for instance in the case of the epidemic of the Nipah Virus between 2001 and 2007 [6]. With regards to the Ebola virus, some twenty outbreaks have been recorded since the discovery of the virus in 1976 [7]. This number of outbreaks undoubtedly underestimates the total number of emergences because not all emergences necessarily lead to the spread of the infection from an animal reservoir to the host population [8]. While the reservoir has an important role for causing the emergence of outbreaks, the role of spillover transmission on the incidental epidemiological dynamics is rarely discussed.

**Table 1:** The reservoir is mostly used as defined by the Centre for Disease Control and prevention (CDC). Two other definitions have been proposed to clarify and complete the notion of reservoir in the case of zoonotic pathogens. On the one hand, Haydon et al. (2002) define the reservoir from a practical point of view in order to take into account all hosts epidemiologically connected to the host of interest (i.e. target host), to implement better control strategies. On the other hand, Ashford (1997) establishes a more generalizable definition: for a given pathogen there is a single reservoir. We will use Ashford’s definition, i.e. a model where a pathogen is persistent in the environment of the incidental host.

| Definitions of a reservoir | Authors | Refs |
| --- | --- | --- |
| “any animal, person, plant, soil, substance or combination of any of these in which the infectious agent normally lives” | CDC | [9] |
| “all hosts, incidental or not, that contribute to the transmission to the target host (i.e. the population of interest) in which the pathogen can be permanently maintained” | Haydon et al. | [10] |
| “an ecologic system in which an infectious agent survives indefinitely” | Ashford | [11, 12] |

Theoretically, pathogen spillover is often neglected and it is generaly assumed that the epidemiological dynamics of outbreaks is driven by the ability of the pathogen to propagate within hosts. For instance, a classification scheme for pathogens has been proposed by Wolfe et al. (2007), including five evolutionary stages in which the pathogen may evolve ranging from exclusive animal infection (Stage I) to exclusive human infection (Stage V) (Figure 1) [13]. The intermediate stages are those found for the zoonotic pathogens (Stages II-IV). Lloyd-Smith et al. (2009), propose to enhance the classification scheme by differentiating the Stages II-IV by ability of pathogens to propagate between individuals in the incidental host (i.e. as a function of the basic reproductive ratio *R*_0_): the non-contagious pathogens (*R*0 = 0, Stage II), pathogens barely contagious inducing stuttering chains of transmission (0 < *R*_0_ < 1, Stage III) and contagious pathogens inducing large outbreaks (*R*_0_ > 1, Stage IV) [14]. However, the role of the reservoir is not clearly defined, and spillover effects on the epidemiological dynamics are not discussed.

**Figure 1:**
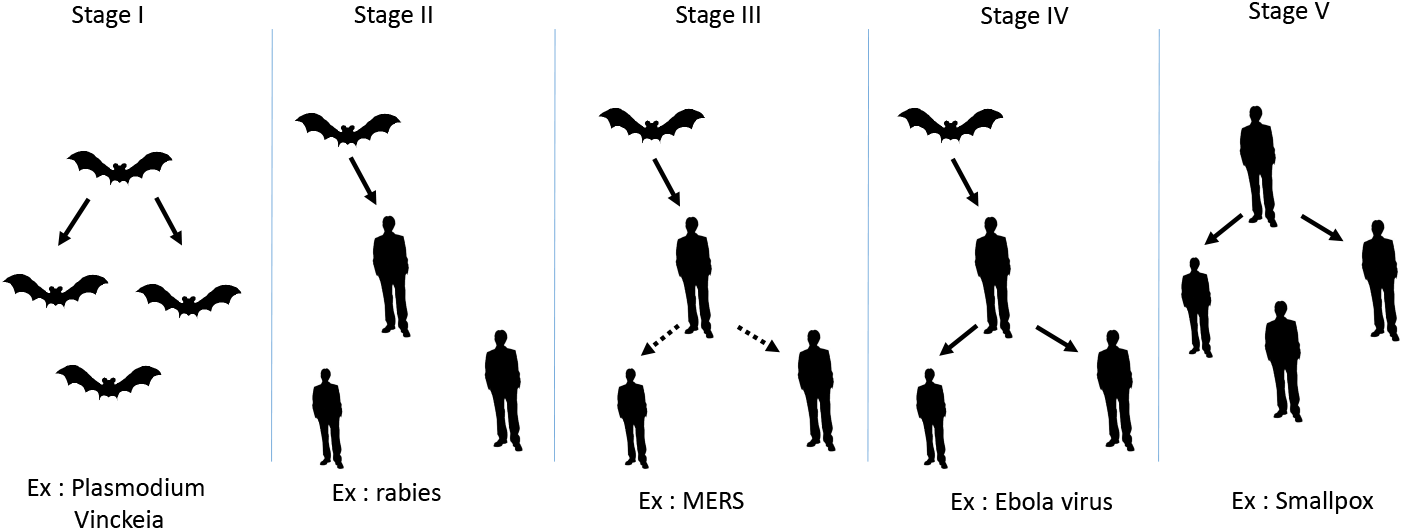
Representation of the evolutionnary stages proposed by Wolfe et al. (2007) in which a pathogen may evolve from infecting only animals to infecting only humans. Each stage corresponds to a specific epidemiological dynamics in the incidental host. Stage II corresponds to few spillovers from animals (e.g. bats) to humans with no possible transmission between humans. Stage III corresponds to few stuttering chains of transmission between humans that go extinct (no outbreaks). Stage IV corresponds to large outbreaks in human population but the pathogen cannot be maintained without the reservoir.

Only a few models have investigated the dynamics of EIDs by taking into account explicitly the transmission from the reservoir to the incidental host. Lloyd-Smith et al. (2009) have analysed 442 modelling studies of zoonotic pathogens and concluded that models incorporating spillover transmission are dismayingly rare [14]. More recent models aimed at investigating the dynamics of EIDs by taking into account the spread of the pathogen using multi-hosts processes but disregarding the persistence of the pathogen in the reservoir [15], or by focusing on the dynamics and conditions of persistence of the pathogen between two populations [16]. Others have considered an endemic reservoir but those models are disease-specific and do not generate generalizable dynamics [17, 18]. More recently, Singh and Myers (2014) developed a Susceptible-Infected-Recovered (SIR) stochastic model coupled with a constant force of infection. Such model is mostly interested in the effect of population size and its impact on the size of an outbreak [19]. However, this approach does not allow teasing apart the contribution of incidental host transmission from that of the transmission from the reservoir in modulating the dynamics of zoonotic pathogens.

In this paper, we aim to provide general insights into the dynamics of a zoonotic pathogen (i.e. pathogens classified in stages II-IV) emerging from a reservoir and its ability to propagate in an incidental host. To do so, we developed a theoretical model that can dissociate the effect of between-host (i.e. direct) transmission from the effect of spillover (i.e. reservoir) transmission. A multi-hosts process with a reservoir and an incidental host are considered. The epidemiological processes are stochastic, which is particularly relevant in the case of transmission from the reservoir and more realistic. The model makes a number of assumptions. First, the epidemiological processes are much faster than the demographic processes. Second, the pathogen in the reservoir is considered as endemic and might contaminate recurrently the incidental host. Third, an individual cannot become susceptible after having been infected. As a consequence, the total number of susceptible individuals in the incidental host decreases during the epidemic. This is what is expected for an epidemic spreading locally during a short period of time (at the scale of a few thousands individuals during weeks or months, depending on the disease and populations considered). We then harness the model to predict the effects of both spillover transmission and direct transmission on the number and the size of outbreaks. Outbreaks occur when the number of cases of disease increases above what would normally be expected. We show that the recurrent emergence of the pathogen from the reservoir in the incidental host is as important as the transmission between individuals of the incidental host. We discuss the implications of these results for the classification of pathogens proposed by Lloyd-Smith et al. (2009).

## 2. Model

A Stochastic Susceptible-Infected-Recovered (SIR) compartmental transmission model [20] with recurrent introduction of the infection into an incidental host by a reservoir is considered (Figure 2). Our goal here is not to study a disease in particular but to provide general insights of the reservoir effect on the epidemiological dynamics of the incidental host. The infection is assumed to propagate quickly relatively to other processes such as pathogen evolution and demographic processes. The reservoir is defined as a compartment where the pathogen is persistently maintained, this pathogen is then considered as endemic. The population is fully mixed. An individual can be infected through two ways of transmission, from the reservoir by the spillover transmission and by direct contact between individuals. We neglect reverse infection from the incidental host to the reservoir.

**Figure 2:**
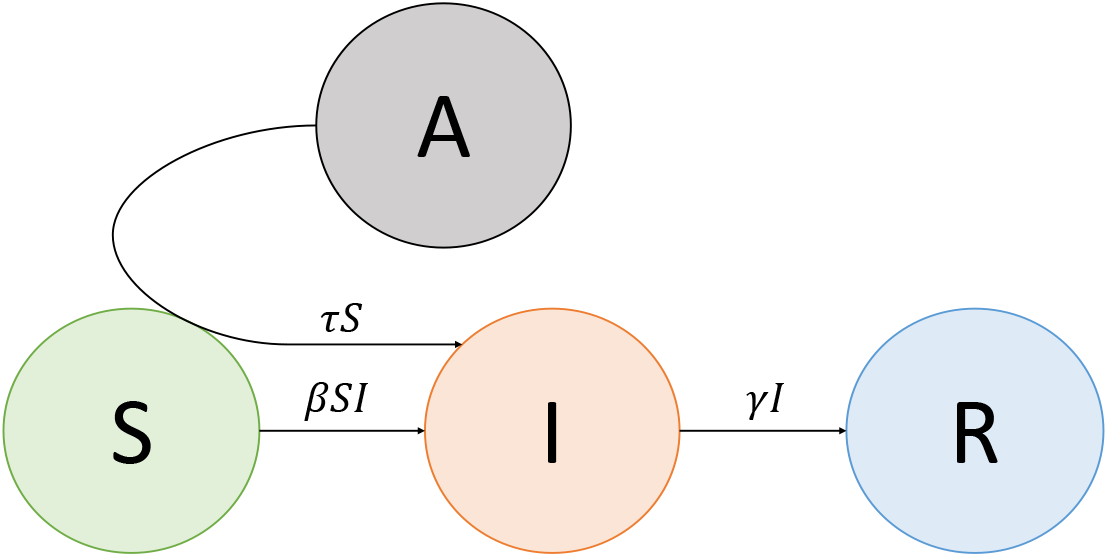
Representation of the stochastic model with transition rates. A reservoir (*A*) has been added to a classical SIR model where the pathogen is persistent. Individuals are characterized by their epidemiological status in the incidental host (*S:* suceptible; *I*: infected; *R:* recovered). A susceptible individual becomes infected through the transmission by contact at rate *βSI* or through the reservoir at rate *τS.* An infected individual recovers at rate *γ*. Stochastic simulations are performed with the following values: the transmission by contact, expressed as the basic reproductive ratio, 0 < *R*_0_ < 10, the spillover transmission, 10^−1^ < *τ* < 10^−10^ and the recovery rate *γ* = 0.1.

The incidental host is composed of *N* individuals. The infection can spillover by contact between the reservoir and the incidental host at rate *τ S* where *S* is the number of susceptible individuals and *τ* is the rate at which an individual becomes infected from the reservoir. In the incidental host, the infection can propagate by direct contact at rate *βSI* where *I* is the number of infected individuals and *β* is the individual rate of infection transmission. An infected individual can recover at rate *γ*. The direct transmission is expressed in terms of the basic reproductive ratio of the pathogen, *R*_0_, which is widely used in epidemiology. *R*_0_ corresponds to the average number of secondary infections produced by an infected individual in an otherwise susceptible population. The interest of this ratio is mostly the notion of threshold: in a deterministic model, for a pathogen to invade the population, *R*_0_ must be larger than 1 in the absence of reservoir. In a stochastic model, the higher the *R*_0_ the higher the probability for the pathogen to invade the population. In a SIR model, the basic reproductive ratio *R*_0_ equals to 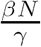. Individuals in the recovered compartment do not contribute anymore to the transmission process. Since we assume that demo-graphic processes are slower than epidemic processes, the number of susceptible individuals decreases during the epidemic due to the consumption of susceptibles by the infection until the extinction of the population.

To analyse the dynamics in the incidental host, three statistics will be studied (i) the mean number of outbreaks, (ii) the mean size of the recurrent outbreaks during an epidemic and (iii) the mean size of the largest outbreak occurring during an epidemic. We consider the appearance of an outbreak when the incidence of the infection exceeds the threshold c and define the maximum size of an outbreak as the largest number of infected individuals during the largest outbreak.

### 2.1. Analysis of the model

#### Stochastic simulations

We performed stochastic individual-based simulations of the epidemics with spillover transmission, using rates as presented in Figure 2, 10 000 simulations are performed for each parameter set. The incidental host is initially (*t* = 0) composed of 1000 susceptible individuals (*N = S =* 1000). The infection is considered as endemic in the reservoir. Simulations are stopped when there are no susceptible individual anymore. We define an outbreak as an excursion (i.e. the stochastic path followed by the population between the first infection to the extinction of the epidemic) during which the number of infected individuals exceeds or equals to the epidemiological threshold *c* (*c* = 5 in the simulations).

#### Approximation by a branching process

The epidemiological model with recurrent introduction of the infection into an incidental host by a reservoir can be approximated by a branching process with immigration from the reservoir to the incidental host at the beginning of the infectious process (thus assuming that individual “birth and death rates of infected individuals” are constant during the starting phase of an outbreak). The individual birth and death rates are respectively *βS,* the transmission rate and *γ*, the recovery rate and the immigration rate corresponds to the spillover rate *τN* at the beginning of the infection. In other words we assume that the number of susceptibles is *N* to study the beginning of the infection, which is a good approximation as long as few individuals have been infected. We distinguish between two regimes in the incidental host, the subcritical regime when *R*_0_ < 1 and the supercritical regime when *R*_0_ > 1. We suppose that at time *t =* 0 a single individual is infected by the spillover transmission.

## 3. Results

### 3.1. The epidemiological dynamics in the incidental host

As illustrated in Figure 3, three patterns are observed (i) a stuttering chain of transmission that goes extinct, i.e. infection spreads inefficiently, (corresponding to Stage II in Wolfe’s classification, see Figure 1), (ii) a large outbreak and few stuttering chains of transmission (corresponding to Stage III, see Figure 1) and (iii) a single large outbreak consuming a large number of susceptible individuals (corresponding to Stage IV see Figure 1).

**Figure 3:**
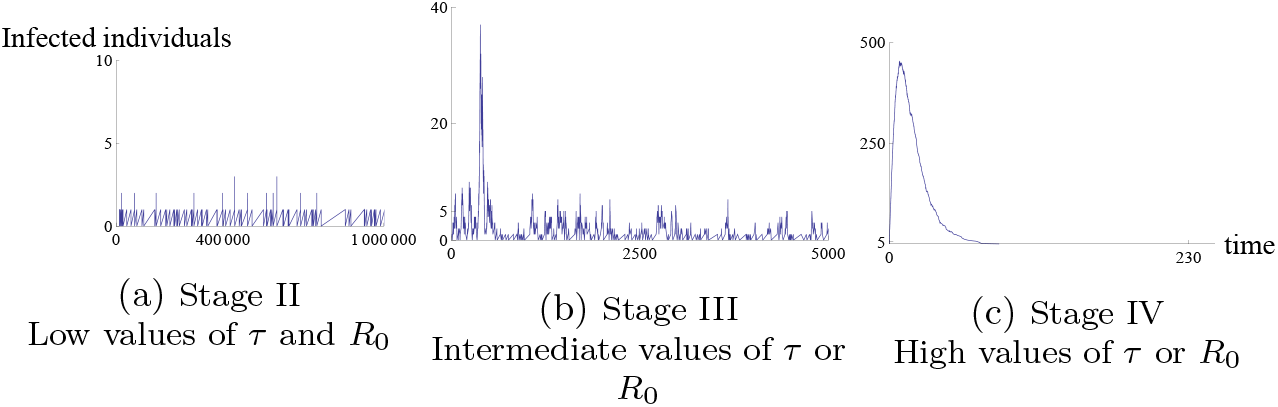
Three examples of epidemiological dynamics corresponding to the three Stages II, III and IV respectively with low values of both direct and spillover transmissions (*R*_0_ = 0.2 and *τ* = 10^−7^), intermediate values of direct or spillover transmission (*R*_0_ = 1.5 and *τ* = 10^−4^), high value of direct or spillover transmission (*R*_0_ = 2 and *τ* = 10^−1^).

Figure 4 shows the roles of *R*_0_ and *τ* in the occurrence of the patterns. Stuttering chains of transmission occur when the pathogen is barely contagious between individuals (small *R*_0_) and when the recurrent emergence of the pathogen (*τ*) is low. At the opposite, when the pathogen is highly contagious (large *R*_0_) or when the spillover transmission is high (*τ*), only one large outbreak is observed. For intermediate values, both dynamics (a large outbreak and stuttering chains of transmission) are observed. The dynamics observed in the three stages depend both of the value of the direct transmission (*R*_0_) and the effect of the reservoir (*τ*).

**Figure 4:**
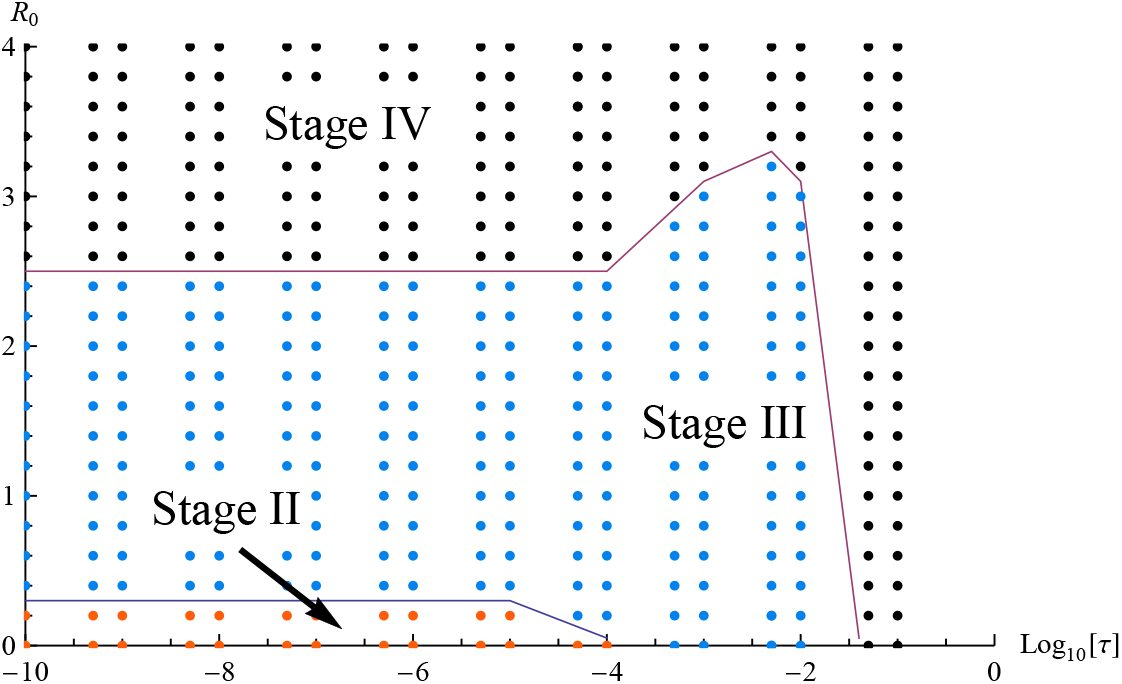
The general epidemiological dynamics is depicted as a function of the direct transmission (*R*_0_) and the spillover transmission (*τ*). The epidemiological dynamics of stochastic simulations are classified following the stages described by Wolfe et al. (2007), Stage II: stuttering chains of transmission (i.e. less than one outbreak), Stage III: one large outbreak and stuttering chains of transmission (i.e. more than one outbreak) and Stage IV: a single large outbreak consuming a large number of susceptible individuals.

### 3.2. What is the effect of the direct transmission on the number of outbreaks when the effect of the reservoir is low?

#### 3.2.1. Case of a barely contagious pathogen (*R*_0_ < 1)

We aim at approximating the mean number of outbreaks in the case where the spillover transmission rate *τ* and the reproductive number *R*_0_ are small (subcritical case corresponding to *R*_0_ < 1). The method of approximation is the following: let us denote by *S*_*i*_ the number of susceptible individuals at the beginning of the *i*-th excursion. During the *i*-th excursion, we set this number of susceptibles to its initial value *S*_*i*_, and consider that the rate of new infections is *βIS*_*i*_. We thus obtain a branching process with individual birth (infection) rate *βS*_*i*_ and individual death (recovery) rate *γ*. When there is no more infected individuals, we compute the mean number of recovered individuals produced by this branching process excursion, denoted by 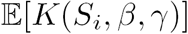, and make the approximation that

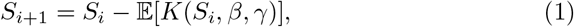

where 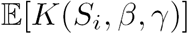 can be computed and equals (see Appendix A):

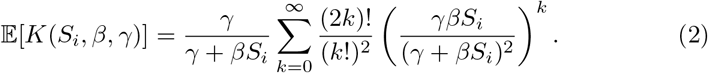

In other words, the initial number of susceptible individuals for the *i* + 1-th excursion is the initial number of susceptible individuals for the *i*-th excursion minus the mean number of recovered individuals produced during the *i*-th excursion under our branching process approximation. We repeat the procedure for the *i* + 1-th excursion, and so on, until *k* satisfies *S*_*k*_ > 0 and *S*_*k*+1_ ≤ 0 (no susceptible anymore). In order to be considered as an outbreak, an excursion has to exceed *c* individuals, where we recall that *c* is the epidemiological threshold. Under our branching process approximation, the probability for the *i*-th excursion to reach the epidemiological threshold (see Appendix A) is:

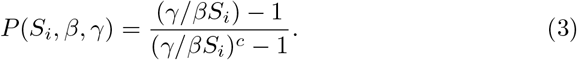

As a consequence, our approximation of the mean number of outbreaks 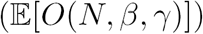 reads:

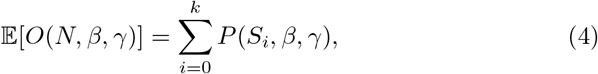

where *S*_1_ = *N*, and the *S*_*i*_’*s* are computed as described in (1).

When the spillover transmission is fixed and small, the mean number of outbreaks computed with the branching process is a good approximation compared to numerical simulations (Figure 5). The spillover transmission added in our model introduces the infection recurrently and allows the infection to spread even for a pathogen barely contagious (*R*0 < 1). According to Figure 5 when *R*_0_ < 1 the number of outbreaks increases when the direct transmission between individuals increases. Indeed, the higher the transmission, the higher the probability for the excursions to reach the epidemiological threshold (*c*). The number of outbreaks can be high because when the direct transmission is smaller than 1, the infection spreads inefficiently and does not consume a large number of susceptibles allowing the next excursion to exceed the epidemiogical threshold.

**Figure 5:**
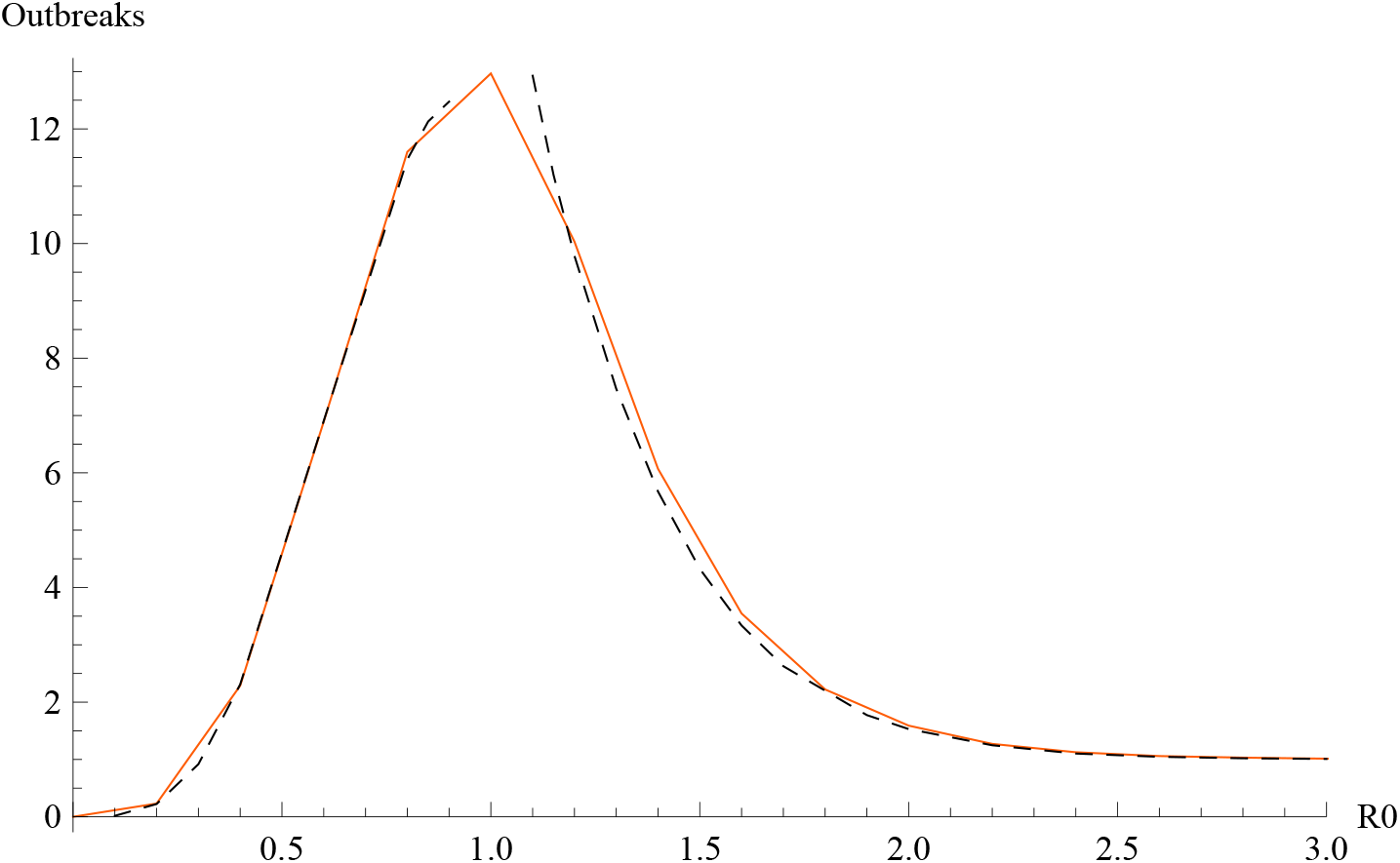
The number of outbreaks is evaluated when the spillover transmission *τ* is small. For the numerical simulations, *τ* = 10^−10^ has been chosen. The orange and black dotted curves represent respectively the numerical simulations and the branching process approximation.

Figure 6 shows that the number of outbreaks is a non-monotonic function of the direct transmission (*R*_0_) and the spillover transmission (*τ*). For intermediate and low values of spillover transmission (10^−6^ < *τ* < 10^−4^), the number of outbreaks increases until *R*0 ~ 1 then decreases. Moreover, we observed an increasing number of outbreaks with *τ* when the pathogen is barely contagious until *τ* is intermediate then decreases when *τ* becomes large 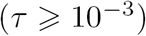.

**Figure 6:**
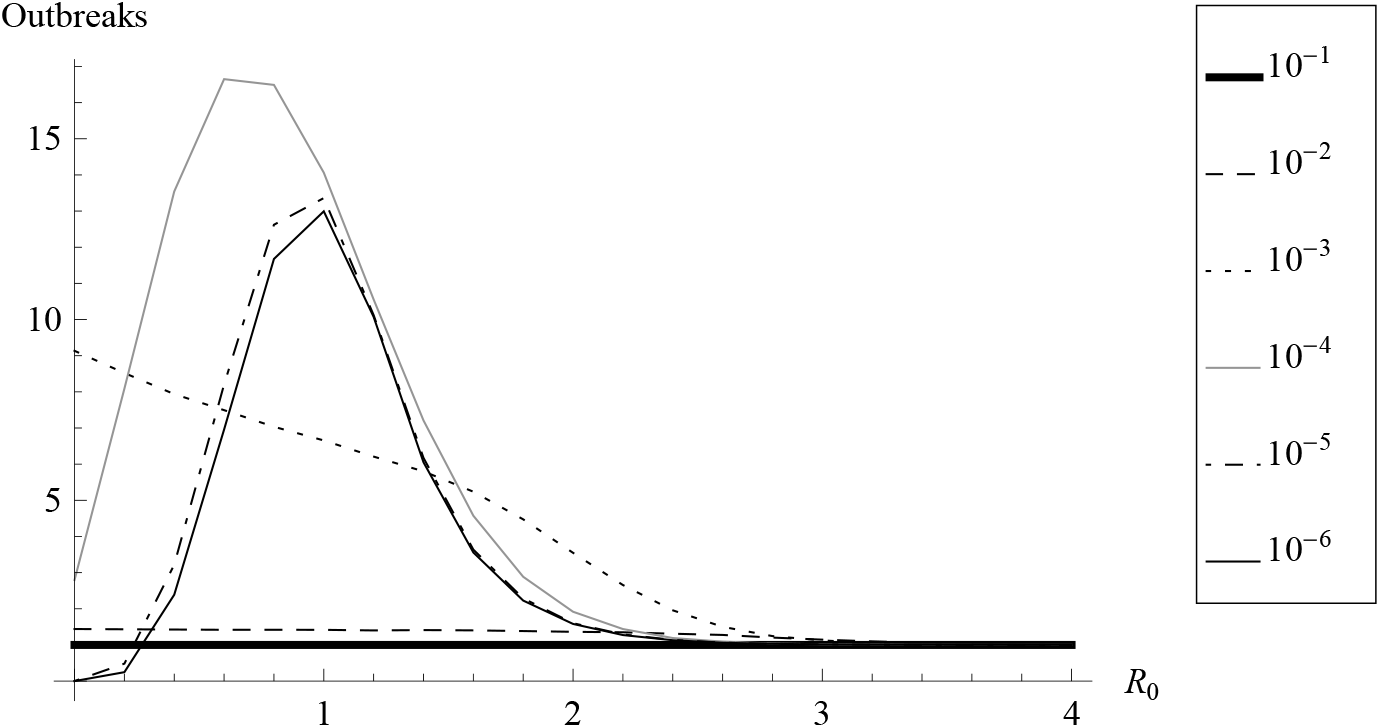
The number of outbreaks as a function of the spillover transmission (10^−6^ < *τ* < 10^−1^). The direct transmission is equal to 0.4.

#### 3.2.2. Case of a contagious pathogen R_0_ > 1

The supercritical case (*R*_0_ > 1) is now considered and the spillover transmission rate (*τ*) is still supposed small.

In this case, two different types of excursions occur in the incidental host: (i) a large outbreak which consumes, with a great probability, a large proportion of susceptible individuals and (ii) multiple excursions before and after the large outbreak. We note *O*_*before*_(*N, β*, *γ*) and *O*_*after*_(*N, β*, *γ*) the number of outbreaks occurring respectively before and after the large outbreak. Because *R*_0_ > 1, the probability to have one large outbreak is high. Hence we make the approximation that one large outbreak occurs during the epidemic, and the total number of outbreaks (*O*_*total*_(*N, β, γ*)) can be approximated by:

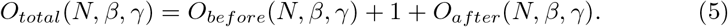

To be part of outbreaks occurring before the large one, an excursion has to satisfy two conditions (i) to have a size higher than the epidemiological treshold *c*, and (ii) to be of a size not too large otherwise it would correspond to the large outbreak. To be more precise, this condition will correspond to the fact that the supercritical branching process used to approximate this excursion does not go to infinity. As a consequence, *O*_*before*_(*N, β, γ*) can be approximated by (See Appendix B):

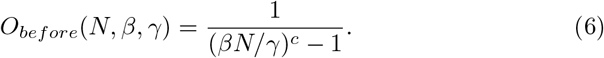

To approximate the number of outbreaks after the large outbreak (*O*_*after*_(*N, β,γ*)), we need to know how many susceptible individuals remain in the population. The number of susceptibles consumed before the large outbreak is negligible with respect to the number of susceptibles consumed during the large outbreak. Hence we can consider that the initial state of the large outbreak is *N* susceptibles, one infected individual and no recovered individual. The number of susceptibles remaining after the large outbreak can be approximated with the deterministic SIR model.

The large outbreak stops when there is no infected individual anymore in the incidental host. Using that 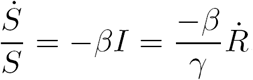, we get:

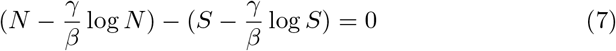

which has one trivial solution (*S*_*i*_ = *N*) and a non-trivial solution with no explicit expression denoted *N*_*after*_(*N, β, γ*). After the large outbreak, the epidemiological threshold for the next excursions, denoted 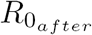, is subcritical (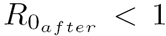) (see Appendix B) and the number of outbreaks after the large one, denoted *O*_*after*_, can be approximated using Equations (2) to (4).

The branching process approximations of the mean number of outbreaks in the supercritical regime, depicted in Figure 5, are close to the mean number of outbreaks found by numerical simulations when the recurrent infection from the reservoir is small. The number of outbreaks decreases when the pathogen becomes highly contagious to reach one outbreak when *R*_0_ > 2.5. When the infection is introduced in the incidental host by the spillover transmission, the probability to reach the epidemiological threshold depends on the direct transmission between individuals. When the direct transmission increases the infection spreads more efficiently consuming a large number of susceptible individuals allowing little or no other excursion to reach the epidemiological threshold and producing only one outbreak when *R*_0_ > 2.5.

### 3.3. What is the effect of the reservoir on the number of outbreaks?

We now focus on the effect of the spillover transmission with a pathogen barely contagious (*R*_0_ < 1) on the number of outbreaks. Because we consider the subcritical case (*R*_0_ < 1), the excursions are small and at the beginning of the epidemiological dynamics, we make the approximation that the spillover transmission rate is constant equal to *τN,* and the direct transmission rate is equal to *βNI.* Using Equation (8) in Singh and Meyer (2014), we get the mean number of infections by the reservoir (*m*) during an excursion under this branching process approximation [19]:

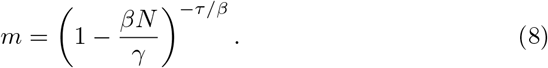

We know that the probability for an excursion to reach the epidemiological threshold *c* in order to be considered as an outbreak is (recall Equation (3) see details in Appendix A):

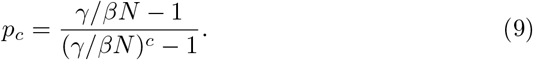

Because the subcritical case is considered, the probability *p*_*c*_ is small, thus allowing the approximation of the probability that the excursion is not an outbreak (See Appendix C for details):

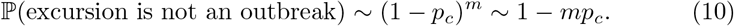

We deduce the probability for the excursion to reach the threshold *c*, (See Appendix C),

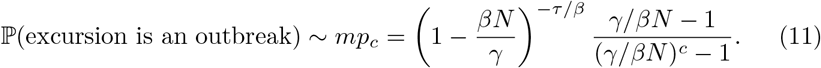

Now, we can study some limit cases of *mp*_*c*_ to approximate the number of outbreaks and understand the effect of the spillover transmission.

When *mp*_*c*_ is small, then the probability to have an excursion reaching the epidemiological threshold *c* is small and the number of outbreaks will be small. *m* small corresponds to a small number of spillovers from the reservoir. According to Figure 7a, when a small effect of spillover transmission (*τ* < 10^−6^) and a small infection reproductive ratio (*R*_0_ < 0.8) are considered then the number of outbreaks is small. In the case of a slightly higher direct transmission (*R*_0_ = 0.8) then each spillover has a non negligible probability to become an outbreak that is why the number of outbreaks for a small spillover transmission rate is higher compared to smaller direct transmission.

**Figure 7:**
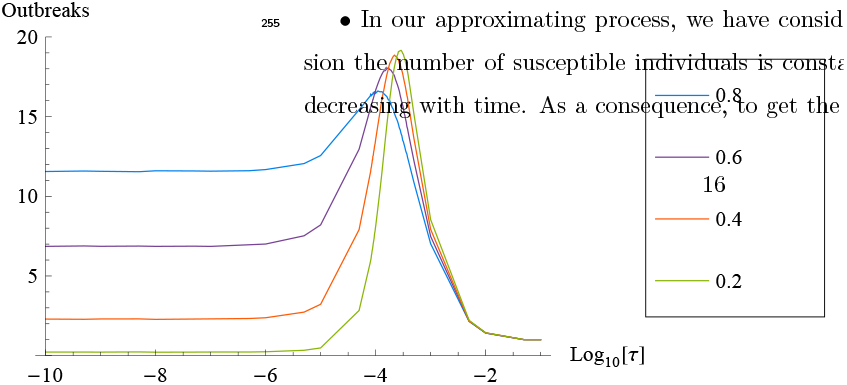
Number of outbreaks as a function of the basic reproductive ratio (0.2 < *R*_0_ < 0:8) and the recurrent emergence rate of a pathogen (*Log*10[10^−10^] < *τ* < *Log*10[10^−1^]) obtained in simulations (lines) and approximations (dots).

When *mp*_*c*_ is large, then the probability that the excursion is not an outbreak, (1 *–p*_*c*_)^*m*^, is small leading to a large outbreak consuming a large number of susceptible individuals. Then few outbreaks will emerge. In Figure 7a, when *τ* is large (*τ* > 10^−2^), that is to say when a large number of spillover transmissions (*m*) arise, only one outbreak is observed because the large number of spillovers prevents the outbreak from dying out.

When *mp*_*c*_ is close to one, each excursion has a non negligible probability to be an outbreak but the number of susceptible individuals consumed is small enough to allow further outbreaks making possible the appearance of multiple outbreaks. We get an approximation of τ for which multiple outbreaks are possible by solving *mp*_*c*_ = 1: in the real process than in the approximation we have done, we need to choose a higher parameter *τ*.

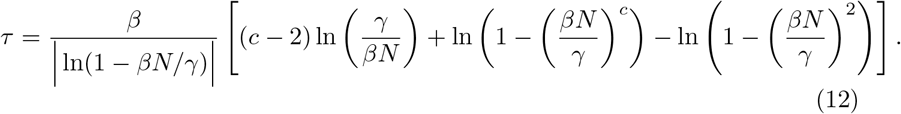

Figure 7b presents the values of *τ* maximising the number of outbreaks and their estimations (dots) obtained by the branching process approximations. First we notice that the branching process approximation gives good results, as the ratio between the real values and their approximations varies between 0.6 and 1.5 for the tested parameter values. Depending on the basic reproductive ratio *R*_0_ we either underestimate (*R*_0_ ≥ 0:6) or overestimate (*R*_0_ ≤ 0:4) the value of *τ* for which the number of outbreaks is maximal due to the choice
of approximating process we have done. Theses errors may come from two approximations made which have variable impacts depending on the value of the direct transmission rate *R*_0_:

- In our approximating process, we have considered that during an excursion the number of susceptible individuals is constant, whereas in reality it is decreasing with time. As a consequence, to get the same number of outbreaks in the real process than in the approximation we have done, we need to choose a higher parameter *τ*.
- We have considered, in our approximating branching process, that when a new individual is infected by the reservoir during an excursion then only one individual is still infected. In reality, there is at least one infected individual but there may be more. As a consequence, the spillover rate may be lower in the real process than in the approximation.

### 3.4 What is the effect of the reservoir on the size of the largest outbreak?

During the epidemic, a large outbreak can occur depending on the value of the direct transmission (*R*_0_) and the spillover transmission (*τ*) and corresponds to the largest number of infected individuals. To analyse the effect of the recurrent emergence of the pathogen on the size of the largest outbreak, we approximate the stochastic model by a SIR deterministic model with a spillover transmission:

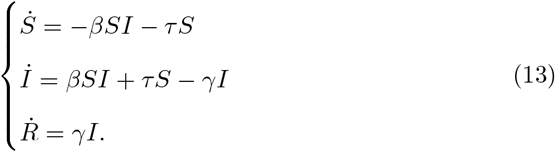

Since no explicit expression of the size of the outbreak can be obtained with the deterministic model, we estimated it using numerical analyses.

Figure 8 shows that the number of infected individuals during the largest outbreak increases with the direct transmission (*R*_0_) and the spillover transmission (*τ*). Furthermore, a large outbreak can even be observed for a pathogen barely contagious (*R*_0_ < 1) when the recurrent emergence of the pathogen is high 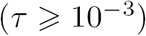.

**Figure 8:**
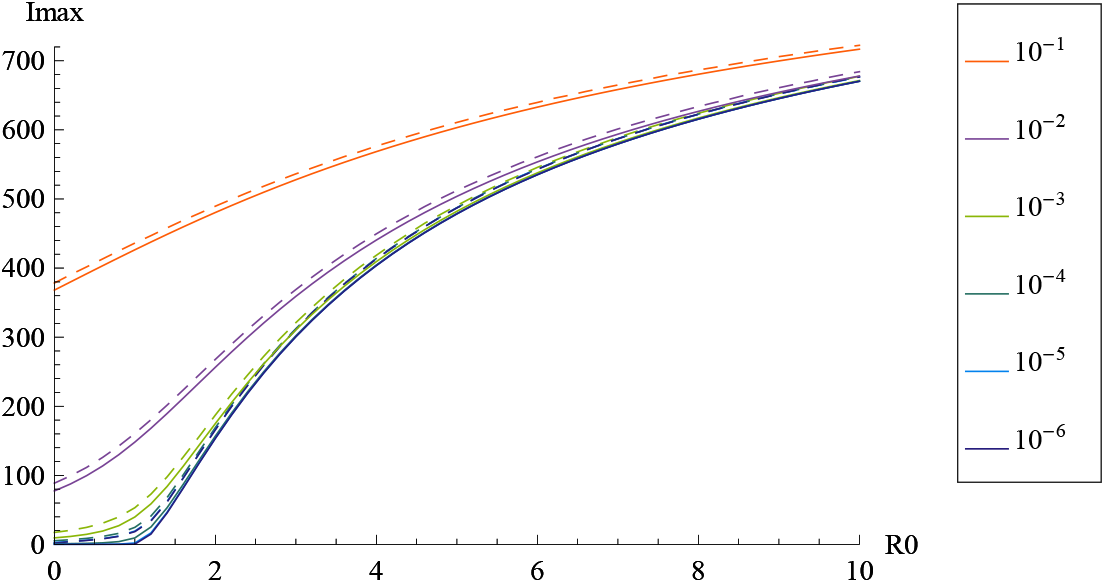
The number of infected individuals during the largest outbreak (*Imax*) is numerically found with the deterministic model (line) and the numerical simulations of the stochastic model (dashed line). The situation is indicated for a reproductive ratio (*R*_0_) varying from 0 to 10 and with the recurrent emergence rate of the pathogen (*τ*) varying from 10^−6^ to 10^−1^.

## 4. Discussion

Zoonotic pathogens constitute one of the most pressing concerns with regards to future emerging diseases, but studies investigating the importance of the role of animal reservoirs for the epidemiological dynamics of infectious diseases are lacking. Indeed, most theoretical works only consider pathogen transmission between conspecifics for predicting disease epidemiology. Here, we build a stochastic SIR model to consider the statistical process underlying a spillover transmission. We draw on the model to predict the number and the size of outbreaks as a function of both the spillover transmission and the direct transmission. The model shows that spillover transmission influences the epidemiological dynamics as much as the transmission by direct contact between individuals. Three different dynamics are observed, ranging from the absence of oubreaks to a single large outbreak; The findings have implications for (1) modelling the dynamics of EIDs, (2) understanding the occurrence of outbreaks in the case of pathogens that are barely contagious and (3) control strategies. The limitations of our approach are discussed.

In our results, the appearance of outbreaks depends on both the transmission from the reservoir and the direct transmission between individuals. Generally, the occurrence of epidemics in humans is attributed to the ability of the pathogen to propagate between individuals. In the case of a single-host process, the notion of the basic reproductive ratio *R*_0_ seems sufficient to evaluate the spread of the pathogen in a population entirely composed of susceptible individuals. In EIDs, *R*_0_ is also used to gauge the risk of pandemics. In this way, Lloyd-Smith et al. (2009) delineate the three stages identified for a zoonotic pathogen [13] by using the ability of the pathogen to spread between individuals. Each stage corresponds to a specific epidemiological dynamics ranging from a non-contagious pathogen making an outbreak impossible (Stage II, *R*_0_ = 0) to a barely contagious pathogen with few outbreaks and stuttering chains of transmission (Stage III, *R*_0_ < 1) to a contagious pathogen making a large outbreak possible (Stage IV, *R*_0_ > 1). The aim of the Wolfe’s classification is to establish each stage in which a zoonotic pathogen may evolve to be adapted to human transmission only, in order to point out pathogens at potential risk of pandemics. However, by taking into account the recurrent emergence of the pathogen from the reservoir, the three dynamics that define the three stages will depend on both the spillover transmission and the direct transmission of the pathogen between individuals. The results suggest that in the case of pathogen spilling recurrently over an incidental host, the direct transmission should not be the only parameter to consider.

The presence of a reservoir and its associated recurrent spillovers dramatically impact the epidemiological dynamics of infectious diseases in the incidental host. Without transmission from the reservoir, the probability to have an outbreak when the pathogen is barely contagious only depends on the direct transmission between individuals, and the outbreak rapidly goes extinct. By contrast, the results show that the recurrent emergence of the pathogen from a reservoir increases the probability to observe an outbreak. Spillover transmission enhances the probability to both observe longer chains of transmission and reach the epidemiological threshold (i.e. threshold from which an outbreak is considered) even for a pathogen barely contagious. Moreover, this coupling model (reservoir-human transmission) allows the appearance of multiple outbreaks depending on both the ability of the pathogen to propagate in the population and the transmission from the reservoir. Zoonotic pathogens such as MERS, Ebola or Nipah are poorly transmitted between individuals [21, 17, 22, 6] yet outbreaks of dozens/hundreds/thousands of infected individuals are observed. We argue that, as suggested by our model, the human epidemic caused by EIDs could be due to recurrent spillover from an animal reservoir.

In the case of zoonotic pathogens, it is of primary importance to distinguish between primary cases (i.e. individuals infected from the reservoir) and secondary cases (i.e. individuals infected from another infected individual) to specify the control strategies to be implemented, in order to optimize the utilization of the public health resources. According to the stochastic SIR model coupled with a reservoir analysed here, the same dynamics can be observed depending on the relative contribution of the transmission from the reservoir and the direct transmission by contact with an infected individual (see Figure 4). For example, a large outbreak is observed either for a high spillover transmission or for a high direct transmission. Empirically, it is generally difficult to distinguish between these two pathways of transmission. Only in the case of non-communicable diseases it is easily possible to measure the importance of the recurrent transmission from the reservoir. Indeed, in this context, all infected individuals originate from a contact with the reservoir. It is the case for the H7N9 virus where most human cases are due to a contact with infected poultry and for which approximately 132 spillovers have been listed during the epidemic of 2013 [23]. For pathogens that are able to propagate from one individual to another, it is difficult to know the origin of the infection, which can be established according to patterns of contacts during the incubation period [6, 17]. Most often, if an infected individual has been in contact with another infected individual in his recent past, direct transmission is considered as the likeliest origin of the infection. However, both individuals might have shared the same environment and thus might have been independently infected by the reservoir. This leads to overestimating the proportion of cases that result from person-to-person transmission. Moreover, when the pathogen infects an individual and the latter does not produce secondary cases then the detection of emergence is unlikely. The proposed stochastic model makes it possible to understand the effects of the infection from the reservoir or from direct transmission on the epidemiological dynamics in an incidental host when empirically this distinction is difficult. Thereafter, the role of control strategies implemented could be evaluated in order to determine the better strategies.

We have considered that the reservoir is a unique population in which the pathogen can persist. The pathogen is then endemic in the reservoir and the reservoir has a constant force of infection on the incidental host. The reservoir can be seen as an ecological system comprising several species or populations in order to maintain the pathogen indefinitely [10]. For example, bat and dromedary camel (*Camelus Dromedarius*) populations are involved in the persistence of MERS-CoV and in the transmission to human populations [24]. In these cases, the assumptions of a constant force of infection can be valid because the pathogen is endemic. However, the zoonotic pathogens can spill over multiple incidental hosts and they can infect each other. In the case of the ebola virus, which infects multiple incidental hosts such as apes, gorillas and monkeys [25], the principal mode of contamination of the human population is non-human primate populations. Moreover, the contact patterns between animals and humans is one of the most important risk factors in the emergence of avian influenza outbreaks [26]. These different epidemiological dynamics with transmission either from the reservoir or from other incidental hosts can largely impact the dynamics of infection observed in the human population, and the investigations of those effects can enhance our understanding of zoonotic pathogens dynamics.

In this paper, we have argued that the conventional way for modelling the epidemiological dynamics of endemic pathogens in an incidental host should be enhanced to account for spillover transmission in addition to conspecifics transmission. We have shown that our stochastic SIR model with a reservoir produce similar dynamics that those found empirically (see the classification scheme for pathogens from [13]). This model can be used to better understand how the ways in which EIDs transmit impact disease dynamics.

## Appendices

In this appendix, we derive results on the branching process approximation stated in Section 3. The main idea of this approximation is the following: when the epidemiological process is subcritical (*R*0 < 1), an excursion will modifiy the state of a small number of individuals with respect to the total population size. During the *i*-th excursions, the direct transmission rate *βSI* will stay close to *βS*_*i*_*I* where *S*_*i*_ denotes the number of susceptibles at the beginning of the *i*-th excursion. Hence, if we are interested in the infected population, the rate *βS*_*i*_*I* can be seen as a constant individual birth rate *βS*_*i*_. Similarly, γ*I*, which is the rate at which an individual in the population recovers, can be interpreted as a constant individual death rate *γ* in the population of infected individuals.

### A. Number of outbreaks in the subcritical case (R_0_ < 1)

In this section, we focus on the number of outbreaks when *R*0 < 1 and when the rate of introduction of the infection by the reservoir is small (τ ≪ 1). That is to say, we consider that each introduction of the infection by the reservoir occurs after the end of the previous excursion created by the previous introduction of the infection by the reservoir. According to Equation (C.1), this approximation is valid as long as the ratio *τ*/*β* is small. We first approximate the mean number of susceptible individuals consumed by an excursion. Let us consider a subcritical branching process with individual birth rate *βS* and individual death rate *γ*. As this process is subcritical, we know that the excursion will die out in a finite time and produce a finite number of individuals. Then from [27], if we denote by *K*[*S, β, γ*] the total number of individuals born during the lifetime of this branching process (counting the initial individual), we know that:

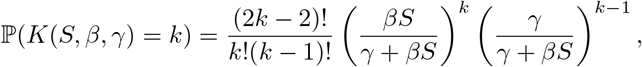

where 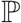 denotes a probability, and hence

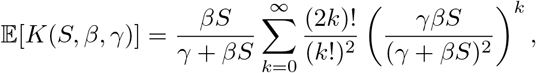

where 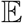 is the expectation. By definition, an excursion is considered as an outbreak only if the maximal number of individuals infected at the same time during this excursion is larger than an epidemiological treshold that we have denoted by *c*. Hence in order to approximate the number of outbreaks we still have to compute the probability for an excursion to be an outbreak. This is a classical result in branching process theory, and can be found in [28] for instance.

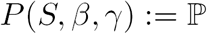(more than *c* individuals infected at the same time) =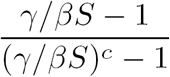

With these results in hands, the method to approximate the mean number of outbreaks is the following. First the probability that the first excursion is an outbreak is

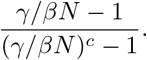

The number of susceptibles at the beginning of the second excursion is approximated by

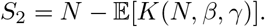

The second excursion has a probability

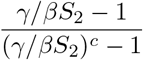

to be an outbreak. The number of susceptibles at the beginning of the third excursion is approximated by

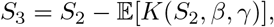

and the third excursion has a probability

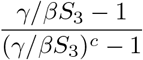

to be an outbreak. And we iterate the procedure as long as there is still a positive number of susceptible individuals. This gives eq. (4).

### B. Number of outbreaks in the supercritical case (R_0_ > 1)

We now focus on the case *R*_0_ = *βN*/*γ* > 1. In this case the approximating branching process will be supercritical and will go to infinity with a positive probability. In the case when the epidemic process describes small excursions, the branching process approximation is still valid, but in the case when it describes a large excursion, then a large fraction of susceptible individuals will be consumed and the branching approximation will not be valid anymore. However, as all the quantities (susceptible, infected and recovered individuals) will be large, a mean field approximation will be a good approximation of the process. Here the mean field approximation will be the deterministic SIR process, whose dynamics is given by:

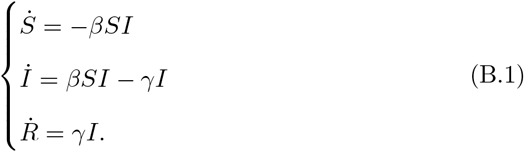

Let us first focus on the small excursions occurring before the large one.

As they are small, they can be approximated by a branching process. Here, unlike in the previous section, the approximating branching process *Z* is supercritical, as *βN* > γ. We can compute its probability to drift to infinity:

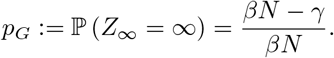

As we will see, a supercritical branching process with individual birth rate *βN* and individual death rate *γ* conditionned to get extinct has the same law as a subcritical branching process with individual birth rate *γ* and individual death rate *βN.* Indeed, if we denote by *Z*_*n*_ the successive values of this branching process, we get for every couple of natural numbers (*n*, *k*):

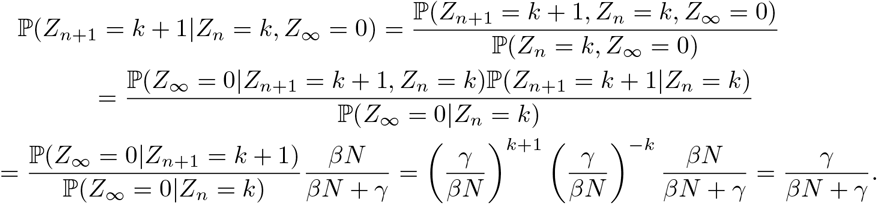

We used again in this series of equalities classical results on branching processes that can be found in [28]. As a consequence, if we denote by *G*[*N, β, γ*] the number of susceptible individuals consumed by the excursion of a supercritical branching process with individual birth rate *βN* and individual death rate *γ* conditionned to get extinct, we get:

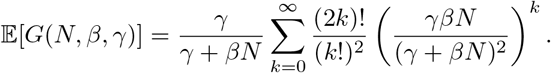

And the probability for this excursion to have a size bigger than the epidemiological treshold *c* is

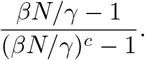

As the number of susceptible individuals stays large until the large excursion occurs, we may keep *N* as the initial number of susceptibles at the beginning of the excursions instead of replacing it by their mean value, as we have done in the previous section.

The different quantities we have just computed allow us to approximate the number of small excursions before the large excursion: in expectation, we have

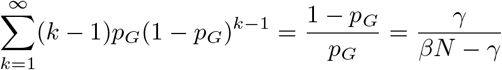

small excursions, which consume

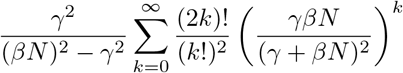

susceptibles and produce

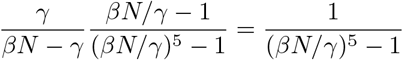

outbreaks.

Now we focus on the large excursion. We will use Equation (B.1) to approximate its dynamics. This equation is well known, and it is easy to obtain the equation satisfied by the final number of susceptible individuals: from (B.1)

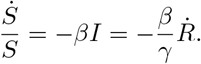

Hence

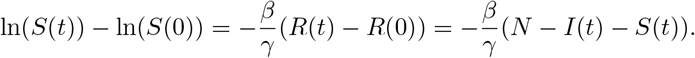

In particular, if we are interested in the time *T*_*f*_ when there is no more infected individual and we suppose that at time 0 there is only one infected individual we get

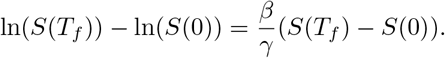

That is to say, *S*(*T*_*f*_) and *S*(0) are related by the equation

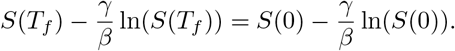

Rigorously, the value of *S*(0) depends on the number of susceptible individuals consumed by the small excursions before the large excursion. But we have seen that this number is small compared to the population size *N*. Hence the number of susceptible individuals remaining after the large excursion can be approximated by the smallest solution of:

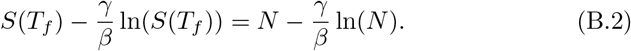

(as the largest solution is *S*(*T*_*f*_) = *N*).

Notice that it is easy to have an idea of the error done for a small variation of the initial state. Indeed, if we denote by *S*_*f*_ the smallest solution of (B.2) and by *S*_*f*_ *– l*(*k*) the solution when *S*(0) = *N* – *k* for a *k* small with respect to *N*, we get:

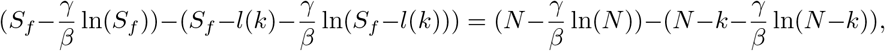

or in other terms

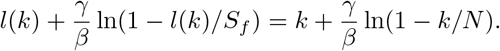

As *k* and *l*(*k*) are small with respect to *N*, this can be approximated by

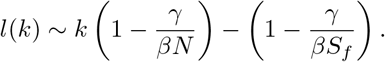

Finally, notice that in (B.1), *S* is a decreasing quantity, and I is a non-negative quantity, which varies continuously. Hence 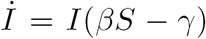 has to be negative before *I* hits 0. As a consequence,

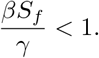

This ensures that the epidemic is subcritical after the large outbreak.

### C. Effect of the reservoir on the number of outbreaks

In this section, we focus on the effect of the reservoir transmission rate (*τ*) on the number of outbreaks when the infection is subcritical (*R*0 < 1). The idea is the following: first, as the excursions of a subcritical branching process are small, we can make the approximation that, at the beginning, the infection rate by the reservoir is constant equal to *τN,* and that the direct transmission rate is equal to *βNI*. Making this approximation allows us to treat the two processes of infection (by contact and by the reservoir) independently. Second, using a result of Singh and Myers (2014), we get the mean number of independent infections by the reservoir during an outbreak under our approximation [19]. As the infections by the reservoir during an outbreak describe a Poisson process, we know that their times are uniformly distributed during the outbreak. As a consequence, we make the approximation that the successive local maxima of the number of infected individuals during the outbreak corresponds to the successive maxima of the outbreaks generated by the independent infections by the reservoir. However, we have to take into account the fact that there is at least one infected individual remaining in the population when a new individual is infected by the reservoir (otherwise the new infected individual would generate a new outbreak). Hence we will make the approximation than the event ‘the excursion does not hit the treshold *c*’ and the event ‘no excursion generated by an infection by the reservoir and with initially two infected individuals, hits the treshold *c*’ are equivalent. Notice that by doing that we underestimate the probability of an excursion to reach any threshold we may explain why we overestimate the spillover transmission *τ* maximizing the number of outbreaks.

Let us denote by m the mean number of infections by the reservoir during an excursion (including the first infection) in our approximation (rate *τN* of infection by the reservoir, and rate *βNI* of direct infections). Then, according to Equation (8) in [19],

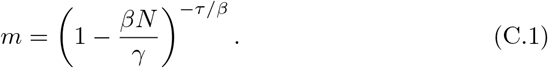

Next, recall that the probability for an excursion of the subcritical branching process with individual birth rate *βN* and individual death rate *γ* and initial state 2 to hit the size *c* is

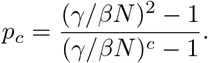

Hence we make the approximation that

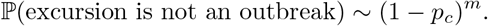

As the approximating branching process is subcritical, the probability *p*_*c*_ is small. Hence, we pursue the approximation by saying

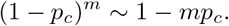

This approximation is valid only if 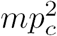 is small, as

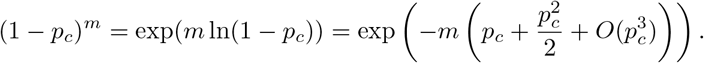

As a consequence, we get

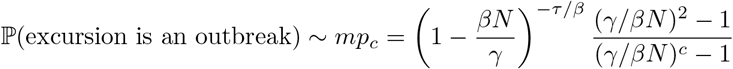

when 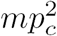 is small. Hence, if *mp*_*c*_ is small, few outbreaks will hit the treshold, and the number of outbreaks will be small. If *mp*_*c*_ is large, (1– *p*_*c*_)^*m*^ is small and the outbreaks will be big, consuming a large number of susceptible individuals. As a consequence, there will be few outbreaks. The case *mp*_*c*_ of order one is the more favorable for outbreaks, as in this case every excursion will have a non negligible probability to be an outbreak, but will be small enough not to consume to many susceptible individuals, allowing other outbreaks to occur. Hence we get an approximation of the optimal *τ* by solving *mp*_*c*_ = 1, which yields:

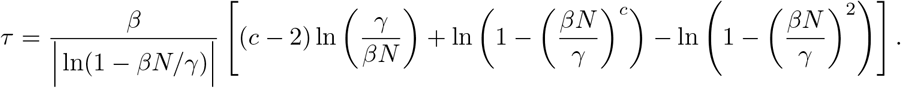

## Acknowledgments

The authors have been supported by the “Chair Modélisation Mathématique et Biodiversité” of Veolia Environnement-Ecole Polytechnique-Museum National d’Histoire Naturelle-Fondation X, France.

## References

[1] K. E. Jones, N. G. Patel, M. A. Levy, A. Storeydard, D. Balk, J. L. Gittleman, P. Daszak, Global trends in emerging infectious diseases, Nature 451 (2008) 990–994. doi:10.1038/nature06536.

[2] L. H. Taylor, S. M. Latham, M. E. J. Woolhouse, Risk factors for human disease emergence, Philosophical Transactions of the Royal Society of London 356 (2001) 983–989. doi:10.1098/rstb.2001.0888.

[3] D. M. Morens, A. S. Fauci, Emerging infectious diseases: Threats to human health and global stability, PloS Pathogens – (7) (2013) e1003467. doi:10.1371/journal.ppat.1003467.

[4] K. A. Murray, P. Daszak, Human ecology in pathogenic landscapes: two hypotheses on how land use change drives viral emergence, Current Opinion in Virology 3 (2013) 79–83. URL http://dx.doi.org/10.1016/j.coviro.2013.01.006

[5] F. Keesing, L. K. Belden, P. Daszak, A. Dobson, D. Harvell, R. D. Holt, P. Hudson, A. Jolles, K. E. Jones, C. E. Mitchell, S. S. Myers, B. T, R. S. Ostfeld, Impacts of biodiversity on the emergence and transmission of infectious diseases, Nature 468 (647) (2010) 647–652. doi:10.1038/nature09575.

[6] S. P. Luby, M. J. Hossain, E. S. Gurley, A. Be-N, S. Banu, S. U. Khan, N. Homaira, P. A. Rota, P. E. Rollin, J. A. Comer, E. Kenah, T. G. Ksiazek, M. Rahman, Recurrent zoonotic transmission of nipah virus into humans, bangladesh, 20012007, Emerging Infectious Diseases 15 (8) (2009) 1229–1235. doi:10.3201/eid1508.081237.

[7] M.-A. de La Vega, D. Stein, G. P. Kobinger, Ebolavirus evolution: Past and present, Plos Pathogens 11 (11).

[8] Z. Jesek, M. Y. Szczeniowski, J. J. Muyembe Tamfum, J. B. McCormick, D. L. Heymann, Ebola, The Journal of Infectious Diseases 179.

[9] Center for disease control and prevention. URL https://www.cdc.gov/hantavirus/resources/glossary.html

[10] D. T. Haydon, S. Cleaveland, L. H. Taylor, Laurenson, Identifying reservoirs of infection: A conceptual and practical challenge, Emerging Infectious Diseases 8 (12) (2002) 1468–1473.

[11] R. W. Ashford, What it takes to be a reservoir, The Belgian Journal of Zoology 127 (1997) 85–90.

[12] R. W. Ashford, When is a reservoir not a reservoir?, Emerging Infectious Diseases 9 (11) (2003) 1495–1496.

[13] N. D. Wolfe, C. P. Dunavan, J. Diamond, Origins of major human infectious diseases, Nature 447 (2007) 279–283. doi:10.1038/nature05775.

[14] J. O. Lloyd-Smith, D. George, K. M. Pepin, V. E. Pitzer, J. R. C. Pulliam, A. P. Dobson, P. J. Hudson, B. T. Grenfell, Epidemic dynamics at the human-animal interface, Science 326 (2009) 1362–1367. doi:10.1126/science.1177345.

[15] S. Singh, D. J. Schneider, C. R. Myers, The structure of infectious disease outbreaks across the animal-human interface, arXiv:1307.4628.

[16] A. Fenton, A. B. Pedersen, Community epidemiology framework for classifying disease threats, Emerging Infectious Diseases 11 (12) (2005) 1815–1821.

[17] G. Chowell, S. Blumberg, L. Simonsen, M. A. Miller, C. Viboud, Synthesizing data and models for the spread of mers-cov, 2013: Key role of index cases and hospital transmission, Epidemics 9 (2014) 40–51. URL http://dx.doi.org/10.1016/j.epidem.2014.09.011

[18] G. T. Nieddu, L. Billings, J. H. Kaufman, R. Forgoston, S. Bianco, Extinction pathways and outbreak vulnerability in a stochastic ebola model, Journal of the Royal Society Interface 14 (2017) 20160847. URL http://dx.doi.org/10.1098/rsif.2016.0847

[19] S. Singh, C. R. Myers, Outbreak statistics and scaling laws for externally driven epidemics, Physical Review 89 (2014) 042108. doi:10.1103/PhysRevE.89.042108.

[20] W. O. Kermack, A. G. McKendrick, A contribution to the mathematical theory of epidemics, Proceedings of the Royal Society of London 115 (772) (1927) 700–721.

[21] A. Zumla, D. S. Hui, S. Perlman, Middle east respiratory syndrome, Lancet 386 (2015) 995–1007. doi:http://dx.doi.org/10.1016/S0140-6736(15)60454-8.

[22] C. L. Althaus, Estimating the reproduction number of ebola virus (ebov) during the 2014 outbreak in west africa, PLOS Currents Outbreaks 6. doi:10.1371/currents.outbreaks.91afb5e0f279e7f29e7056095255b288.

[23] J. Zhou, D. Wang, R. Gao, B. Zhao, J. Song, X. Qi, Y. Zhang, Y. Shi, L. Yang, W. Zhu, T. Bai, K. Qin, Y. Lan, S. Zou, J. Guo, J. Dong, L. Dong, Y. Zhang, H. Wei, W. Li, J. Lu, L. Liu, X. Zhao, X. Li, W. Huang, Biological features of novel avian influenza a (h7n9) virus, Nature 499 (2013) 500–503.

[24] J. S. M. Sabir, T. T. Y. Lam, M. M. M. Ahmed, L. Li, Y. Shen, S. E. M. Abo-Aba, M. I. Qureshi, M. Abu-Zeid, Y. Zhang, M. A. Khiyami, N. S. Alharbi, N. H. Hajrah, M. J. Sabir, M. H. Z. Mutwakil, S. A. Kabli, F. A. S. Alsulaimany, A. Y. Obaid, B. Zhou, D. K. Smith, E. C. Holmes, H. Zhu, Y. Guan, Co-circulation of three camel coronavirus species and recombination of mers-covs in saudi arabia, Science 351 (6268) (2016) 81–84. doi:10.1126/science.aac8608.

[25] H. Ghazanfar, F. Orooj, M. A. Abdullah, A. Ghazanfar, Ebola, the killer virus, Infectious Diseases of Poverty 4 (15). doi:DOI10.1186/s40249-015-0048-y.

[26] A. Meyer, T. X. Dinh, T. V. Nhu, L. T. Pham, S. Newman, T. T. T. Nguyen, D. U. Pfeiffer, T. Vergne, Movement and contact patterns of long-distance free-grazing ducks and avian influenza persistence in vietnam, PloS One doi:https://doi.org/10.1371/journal.pone.0178241.

[27] T. Britton, E. Pardoux, Stochastic epidemics in a homogeneous community, in preparation.

[28] K. B. Athreya, P. E. Ney, Branching processes, Springer, Berlin, Heidelberg, 1972.

